# Utilizing transcriptomics and metabolomics reveal drought tolerance mechanism in *Nicotiana tabacum*

**DOI:** 10.1101/2024.05.06.592846

**Authors:** Quanyu Yin, Zhao Feng, Zhichao Ren, Hui Wang, Dongling Wu, Amit Jaisi, Mengquan Yang

**Affiliations:** National Tobacco Cultivation, Physiology and Biochemistry Research Center, College of Tobacco Science, Henan Agricultural University, Zhengzhou, Henan, China; Luoyang Municipal Tobacco Company, Henan Tobacco Industry Co. LTD, Luoyang, Henan, China; School of Pharmacy, Walailak University, Thasala, Nakhon Si Thammarat, 80160, Thailand; Drug and Cosmetics Excellence Center, Walailak University, Thasala, Nakhon Si Thammarat, 80160, Thailand

**Keywords:** *Nicotiana tabacum*; transcriptome, metabolome, physiological characteristics, drought

## Abstract

The development and growth of plants are significantly impacted by adverse surroundings, particularly drought conditions. The yield and quality of plants, in particular, are heavily reliant on the presence of favorable growth conditions. Here, we performed comprehensive research to investigate phenotype, physiological characteristics, transcriptomic and metabolomic changes in *Nicotiana tabacum* (*N. tabacum*) in responses to drought stress (DS). This work aimed to investigate the detailed responses of *N. tabacum* to DS under different drought conditions (CK, well-watered; LD, light drought; MD, moderate drought and SD, severe drought). *N. tabacum* grew normally under CK but was inhibited under LD, MD and SD stress; the relative water content, transpiration rate and protective enzyme activity significantly influenced under DS. In the LD/CK, MD/CK and SD/CK comparison groups, there were 7483, 15558 and 16876 differentially expressed genes (DEGs), respectively, and 410, 485 and 523 differentially accumulated metabolites (DAMs), respectively. The combined analysis of transcriptomic and metabolomic data unveiled the significant involvement of phenylpropanoid biosynthesis in the *N. tabacum*’s response to drought stress. These findings characterized the key metabolites and genes in responses to drought stress in *N. tabacum*, hence offering valuable insights into the underlying mechanisms driving these responses to DS and maintaining plant health under climate change.

## 1. Introduction

Rising global temperatures and land desertification have reduced soil water retention capacity and moisture content, resulting in decreased crop yields [1-3]. Drought is one of the main factors limiting plant survival and growth, affecting plant growth, development, physiological and biochemical metabolic processes, and ecological environment at all stages [4, 5]. Under the increasingly severe threat of drought worldwide, the adaptation of plants to drought stress and their drought resistance mechanisms have become popular research topics in many fields, such as environmental science, ecology and genetics [6, 7]. Drought stress can alter cell membrane structure, enzyme activity, reactive oxygen species accumulation and production, stomatal control, and other gas exchange processes in plants [8-11]. Therefore, exploring the mechanism of plant tolerance against drought stress is crucial for global food security.

Polyphenols, with over 8000 compounds, are the largest and most investigated plant-specific metabolites, which are biosynthesized through phenylpropanoid biosynthesis [12, 13]. Plants produce polyphenols to carry out essential physiological functions, including the regulation of growth, the synthesis of flavonoids, lignin and pigments. In addition, polyphenols have the ability to remove harmful reactive oxygen species (ROS). Plants exhibit increased polyphenol production as an adaptive response to many abiotic and biotic stresses such as salinity, heavy metals, drought, temperature, ultraviolet radiation, plant pathogens and others [14]. For example, foxtail millet that had adapted to drought conditions showed enhanced ability to withstand drought, as indicated by improved growth, increased antioxidant defense, accumulation of polyphenols, and altered gene expression associated with polyphenol biosynthesis [15]. All phenolic compounds increased in Pinto beans under drought stress [16]. Drought enhanced the content of phenolic compounds, flavonoids, and antioxidant activities in rapeseed, which was associated with higher epicatechin, and caffeic, and syringic acids [17].

Plants respond to drought conditions by activating many signaling pathways, which leads to major alterations in gene expression and the plant metabolome [18-21]. For instance, metabolomics and transcriptomics analysis revealed that severe drought stress promotes alkaloid biosynthesis and asd, DapL, PrAOs, DDC, and ALDH are the key genes regulating alkaloid biosynthesis in *Sophora alopecuroides* [18]. Metabolomics analysis indicated that flavonoids were accumulated under drought stress [19]. Transcriptome and metabolome investigations showed that *Illicium difengpi* increased glutathione, flavonoids, polyamines, soluble sugars, and amino acids during drought stress, improving cell osmotic potential and antioxidant activity. [20]. Drought-tolerant peanuts use an altered terpene skeleton synthesis route to supply primary and secondary metabolism [21].

*N. tabacum* is an economically important crop, which is cultivated worldwide and also used as an excellent model plant and bioreactor for research studies [22, 23]. However, tobacco is susceptible to drought stress. Drought stress has long posed a threat to the effective production of *N. tabacum*, particularly the water deficiency, which has greatly restricted the yielding and quality formation [24-26]. *N. tabacum* provides a valuable platform for investigating the intricate mechanisms underlying responses to drought stress. In this study, transcriptome and metabolome analysis were used to evaluate the physiological characteristics, gene expression and metabolite profiles of *N. tabacum* under drought stress. This research aims to unravel the multifaceted effects of drought on *N. tabacum* at the transcriptomic and metabolomic levels, shedding light on the adaptive strategies employed by plants in the wake of water scarcity. Additionally, it may also inspire the identification of stress-inducible genes for molecular breeding and the development of climate-resilient agricultural practices.

## 2. Materials and methods

### 2.1. Plant materials and drought treatment

*N. tabacum* cultivar K362 was used for all experiments. The seeds were germinated in a 96-well tray in Hongland Nutrient solution (25 °C, 16-h light/8-h dark), and seedlings were transplanted in plastic pots (35 cm in diameter and 40 cm in height, one plant per pot) when the root length of the seedlings grew to approximately 2 cm. All experiments were performed in the greenhouse of Henan Agricultural University (Xuchang, China).

After 22 days, the healthy plants were subjected to drought treatment. 120 plants (30 plants per treatment) were subjected to drought treatment. Seedlings were grown under 4 different conditions: well-watered (CK), light drought (LD), moderate drought (MD) and severe drought (SD). 2 days after drought treatment, the plants were sampled for the first time. And then, the plants were sampled every 5 days. All the samples (harvested at 4 time points) were marked 1st, 2nd, 3rd and 4th, respectively. The plants were harvested and separated into leaves and roots. The samples were quickly frozen in liquid nitrogen and stored at −80 °C for physiological parameters determination and multi-omics analysis.

### 2.2. Detection of relative water content and transpiration rate during drought treatment

The leaf samples were harvested for the relative moisture content measurement [27]. After weighing fresh tobacco leaves, absorb water to saturation and weigh them, then dry and weigh them. Calculate the percentage of fresh leaf water content in saturated moisture content as the relative water content. The transpiration rate was measured by using a Li-6400 portable photosynthetic instrument (Li Cor Inc., USA) from 9:00 AM to 11:30 AM. Origin 2021 was used for plotting.

### 2.3. Protective enzyme activity determination

Freeze the same part of the leaves with liquid nitrogen and store them in a refrigerator at -80 □. The activities of catalase (CAT), peroxidase (POD), superoxide dismutase (SOD) and ascorbic acid peroxidase (APX) were measured using the Solyberg reagent kit.

### 2.4. Transcriptome sequencing and analysis

Total RNA was extracted from the leaves of tea plants using a TRIzol reagent kit according to the manufacturer’s protocol (Invitrogen, CA, USA). After RNA purity and quantification were evaluated using the NanoDrop 2000 spectrophotometer (Thermo Scientific, USA). RNA integrity was assessed using the Agilent 2100 Bioanalyzer (Agilent Technologies, Santa Clara, CA, USA). Then the libraries were constructed using VAHTS Universal V6 RNA-seq Library Prep Kit according to the manufacturer’s instructions. The transcriptome sequencing and analysis were conducted by OE Biotech Co., Ltd. (Shanghai, China).

The libraries were sequenced on an llumina Novaseq 6000 platform and 150 bp paired-end reads were generated. Raw reads for each sample were generated. Raw reads of fastq format were firstly processed using fastp and the low-quality reads were removed to obtain the clean reads [28]. Then clean reads for each sample were retained for subsequent analyses. The clean reads were mapped to the reference genome using HISAT2 [29]. FPKM of each gene was calculated and the read counts of each gene were obtained by HTSeq-count [30, 31]. PCA analysis was performed using R (v 3.2.0) to evaluate the biological duplication of samples.

Differential expression analysis was performed using the DESeq2 [32]. Q value < 0.05 and foldchange > 2 or foldchange < 0.5 were set as the threshold for significantly differential expression genes (DEGs). Hierarchical cluster analysis of DEGs was performed using R (v 3.2.0) to demonstrate the expression pattern of genes in different groups and samples. The radar map of the top 30 genes was drawn to show the expression of up-regulated or down-regulated DEGs using R packet ggradar.

Based on the hypergeometric distribution, GO [33], KEGG pathway [34], Reactome and WikiPathways enrichment analysis of DEGs were performed to screen the significantly enriched term using R (v 3.2.0), respectively. R (v 3.2.0) was used to draw the column diagram, the chord diagram and bubble diagram of the significant enrichment term.

Gene Set Enrichment Analysis (GSEA) was performed using GSEA software [35, 36]. A predefined gene set was used for analysis, and the genes were ranked according to the degree of differential expression in the two types of samples. Then it is tested whether the predefined gene set was enriched at the top or bottom of the ranking list.

### 2.5. Sample preparation and LC-MS analysis

All samples weighing 60 mg each (12 samples, three replicates for each group) were mixed with methanol extract (70%, 0.6 mL, L-2-chlorophenylalanine, succinic acid-d_4_, L-valine-d_8_, Cholic acid-d_4_ (4 μg/mL) as internal standard), and ground in a grinder (60 Hz, 2 min). Then, the samples were vortexed for 30 min and centrifuged at 120,00 rpm for 3 min at 4 °C. After transferring the supernatant to the microporous membrane (pore size: 0.22 μm), the filtered samples were analyzed by the UPLC-MS/MS. Waters ACQUITY UPLC I-Class plus/Thermo QE was used to conduct liquid chromatography-mass spectrometry (LC-MS) analyses.

The metabolomic data analysis was performed by Shanghai Luming biological technology co., LTD (Shanghai, China). An ACQUITY UPLC I-Class plus (Waters

Corporation, Milford, USA) fitted with Q-Exactive mass spectrometer equipped with heated electrospray ionization (ESI) source (Thermo Fisher Scientific, Waltham, MA, USA) was used to analyze the metabolic profiling in both ESI positive and ESI negative ion modes. An ACQUITY UPLC HSS T3 column (1.8 μm, 2.1×100 mm) were employed in both positive and negative modes. The binary gradient elution system consisted of (A) water (containing 0.1 % formic acid, v/v) and (B) acetonitrile (containing 0.1 % formic acid, v/v) and separation was achieved using the following gradient: 0.01 min, 5% B; 2min, 5% B; 4min, 30% B; 8min, 50% B; 10min, 80% B; 14min, 100% B; 15 min, 100% B; 15.1 min, 5% and 16 min, 5%B. The flow rate was 0.35 mL/min and column temperature was 45□. All the samples were kept at 10□ during the analysis. The injection volume was 3 μL.

The mass range was from m/z 100 to 1,000. The resolution was set at 70,000 for the full MS scans and 17500 for HCD MS/MS scans. The Collision energy was set at 10, 20 and 40 eV. The mass spectrometer operated as follows: spray voltage, 3800 V (+) and 3200 V (−); sheath gas flow rate, 35 arbitrary units; auxiliary gas flow rate, 8 arbitrary units; capillary temperature, 320°C; Aux gas heater temperature, 350°C; S-lens RF level, 50.

### 2.6. Metabolomic analysis

The original LC-MS data were processed by the software Progenesis QI V2.3 (Nonlinear, Dynamics, Newcastle, UK) for baseline filtering, peak identification, integral, retention time correction, peak alignment, and normalization. Main parameters of 5 ppm precursor tolerance, 10 ppm product tolerance, and 5% product ion threshold were applied. Compound identification was based on the precise mass-to-charge ratio (M/z), secondary fragments, and isotopic distribution using The Human Metabolome Database (HMDB), Lipidmaps (V2.3), Metlin, and self-built databases. The extracted data were then further processed by removing any peaks with a missing value (ion intensity = 0) in more than 50% in groups, by replacing the zero value by half of the minimum value, and by screening according to the qualitative results of the compound. Compounds with resulting scores below 36 (out of 60) points were also deemed to be inaccurate and removed. A data matrix was combined from the positive and negative ion data.

The matrix was imported in R to carry out Principle Component Analysis (PCA) to observe the overall distribution among the samples and the stability of the whole analysis process. Orthogonal Partial Least-Squares-Discriminant Analysis (OPLS-DA) and Partial Least-Squares-Discriminant Analysis (PLS-DA) were utilized to distinguish the metabolites that differ between groups. To prevent overfitting, 7-fold cross-validation and 200 Response Permutation Testing (RPT) were used to evaluate the quality of the model. Variable Importance of Projection (VIP) values obtained from the OPLS-DA model were used to rank the overall contribution of each variable to group discrimination. A two-tailed Student’s T-test was further used to verify whether the metabolites of difference between groups were significant. Differential metabolites were selected with VIP values greater than 1.0 and p-values less than 0.05.

### 2.7. Combined analysis of genes and metabolites

The pathways that contained both differentially expressed genes (DEGs) and differentially expressed metabolites (DEMs) under the same DS treatment were identified and displayed on the KEGG pathway map. A KGML network was created to visually represent the degree of enrichment for the discovered pathways.

## 3. Results

### 3.1. Phenotypes of *N. tabacum* during DS treatment

As shown **Fig.1**, agronomic traits of *N. tabacum* were influenced a lot after DS-treatment. With the escalation of drought, the plants exhibited evident phenotypic differences, such as severe wilting. Meanwhile, with the period extension of drought treatment, the impact of drought stress on plant phenotypes became more visible. Particularly, during the final phase of sampling time, the phenotypes of the plants exhibited significant variations. Well-watered group (CK) showed greater plant height, higher leaves count, etc. However, severe drought group (SD) had smaller height, lower leaf count, and symptoms of wilting.

**Fig. 1.**
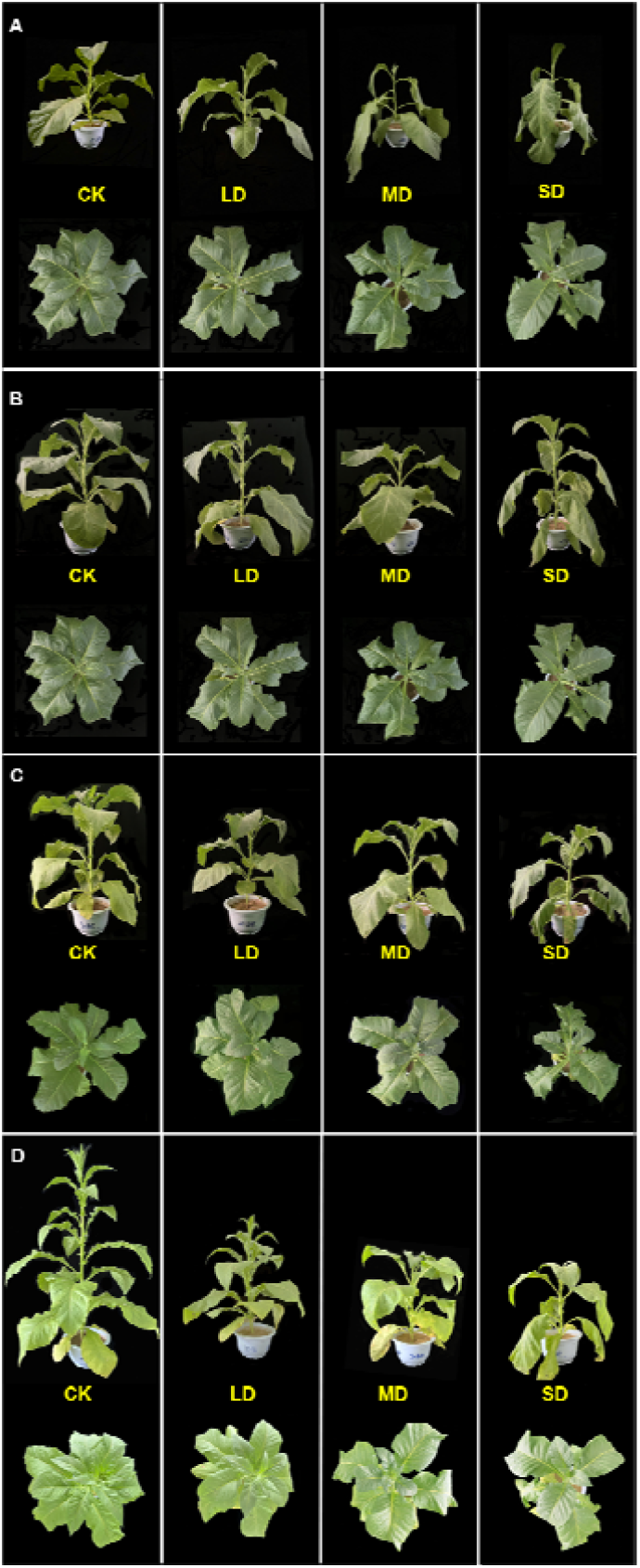
The plant growth status of *N. tabacum* under different drought conditions. A. the first time sampling (1st); B. the second time sampling (2nd); C. the third time sampling (3rd); D. the fourth time sampling (4th). Drought conditions: well-watered (CK); light drought (LD); moderate drought (MD) and severe drought (SD).

Upon analyzing the phenotypic data of the plants, it was observed that drought had a significant impact on both plant height (**Fig.1)**. Significantly, the growth of plants in terms of height exhibited a slow and limited growth during the duration of drought treatments. Conversely, the well-watered plants (CK) displayed the highest growth rate.

### 3.2. Root system development of *N. tabacum* during DS treatment

It is clearly observed that the plant’s root system exhibits robust development when it is well-watered (**Fig.2**). The root system exhibits a tendency to decrease in size as the intensity of the drought stress increases, with the drought stress treatments (LD, MD, and SD) following this pattern in sequential orders. With the extension of drought treatment duration, the severe drought treatment had a significant impact on the growth of the root system, resulting in the most severe impacts at each sampling time.

**Fig. 2.**
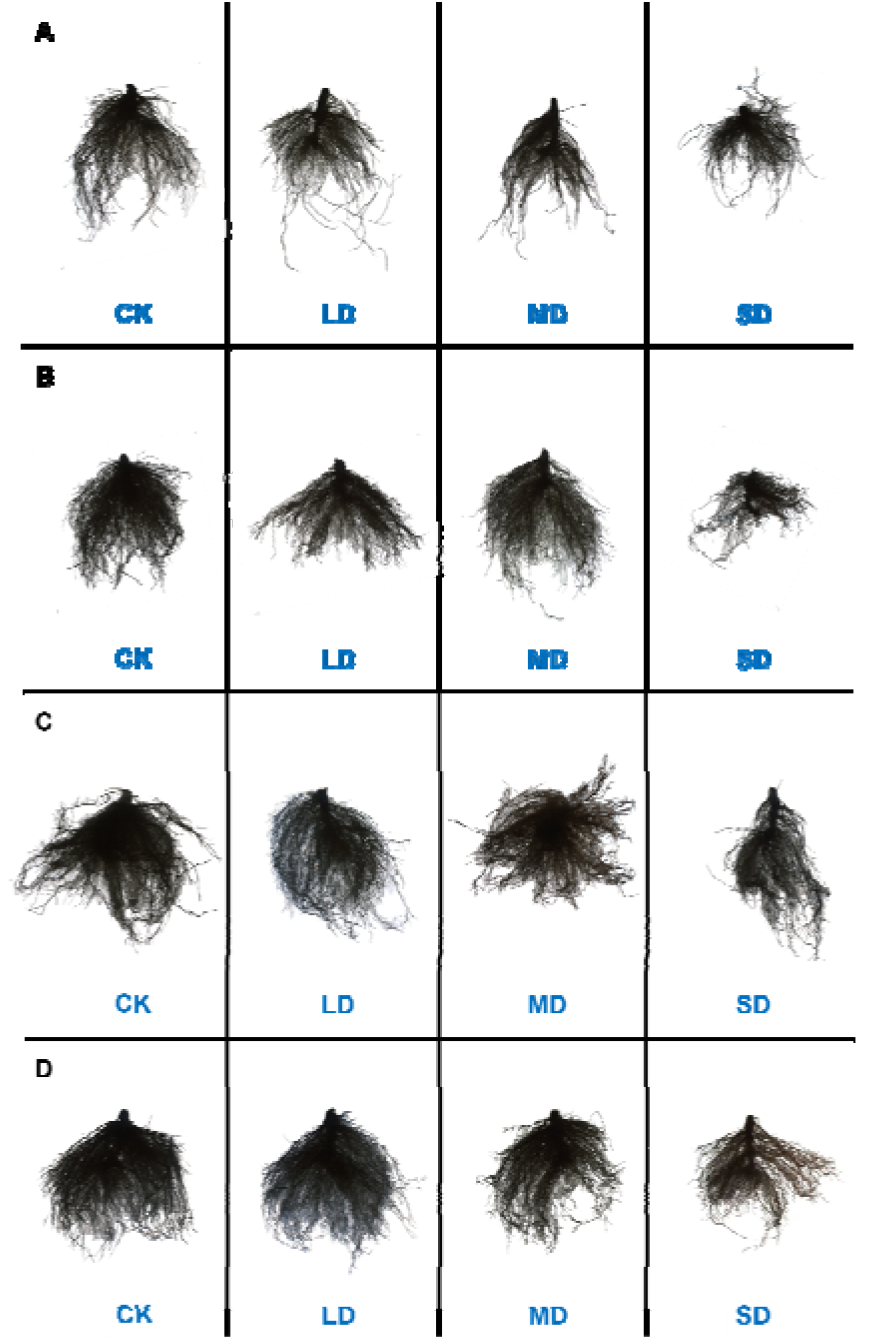
The root system status of *N. tabacum* under different drought conditions. A. the first time sampling (1st); B. the second time sampling (2nd); C. the third time sampling (3rd); D. the fourth time sampling (4th). Drought conditions: well-watered (CK); light drought (LD); moderate drought (MD) and severe drought (SD).

### 3.3. Relative water content, transpiration rate and protective enzyme activity during DS treatment

For the purpose of evaluate the growth status of plants suffering drought stress, we measured the relative water content and transpiration rate of plants at multiple times during the drought treatment. As drought severity increased, there was a little decline in relative water content. As the duration of the drought boosted the amount of water kept on decreasing. Specifically, in the MD and SD treatments, there was a significant reduction in the relative water content at the end of the drought treatment (**Supplementary Fig.1A**). The transpiration rate exhibited a steady decline as the severity of drought increased. Notably, the transpiration rate reached its highest point during the third sampling (**Supplementary Fig.1B**). In addition, the protective enzymes’ activities in *N. tabacum* were measured during the drought treatment. It was discovered that the activities of four endogenous protective enzymes, namely APX (**Fig.3A**), POD (**Fig.3B**), CAT (**Fig.3C**), and SOD (**Fig.3D**), were carried out. The activities of APX, POD, and SOD were all decreased as a consequence of drought treatments (LD, MD, and SD). However, CAT activity showed a tendency to decrease with drought intensification. Meanwhile, the activity of CAT underwent a gradual decrease with the plant growth.

**Fig. 3.**
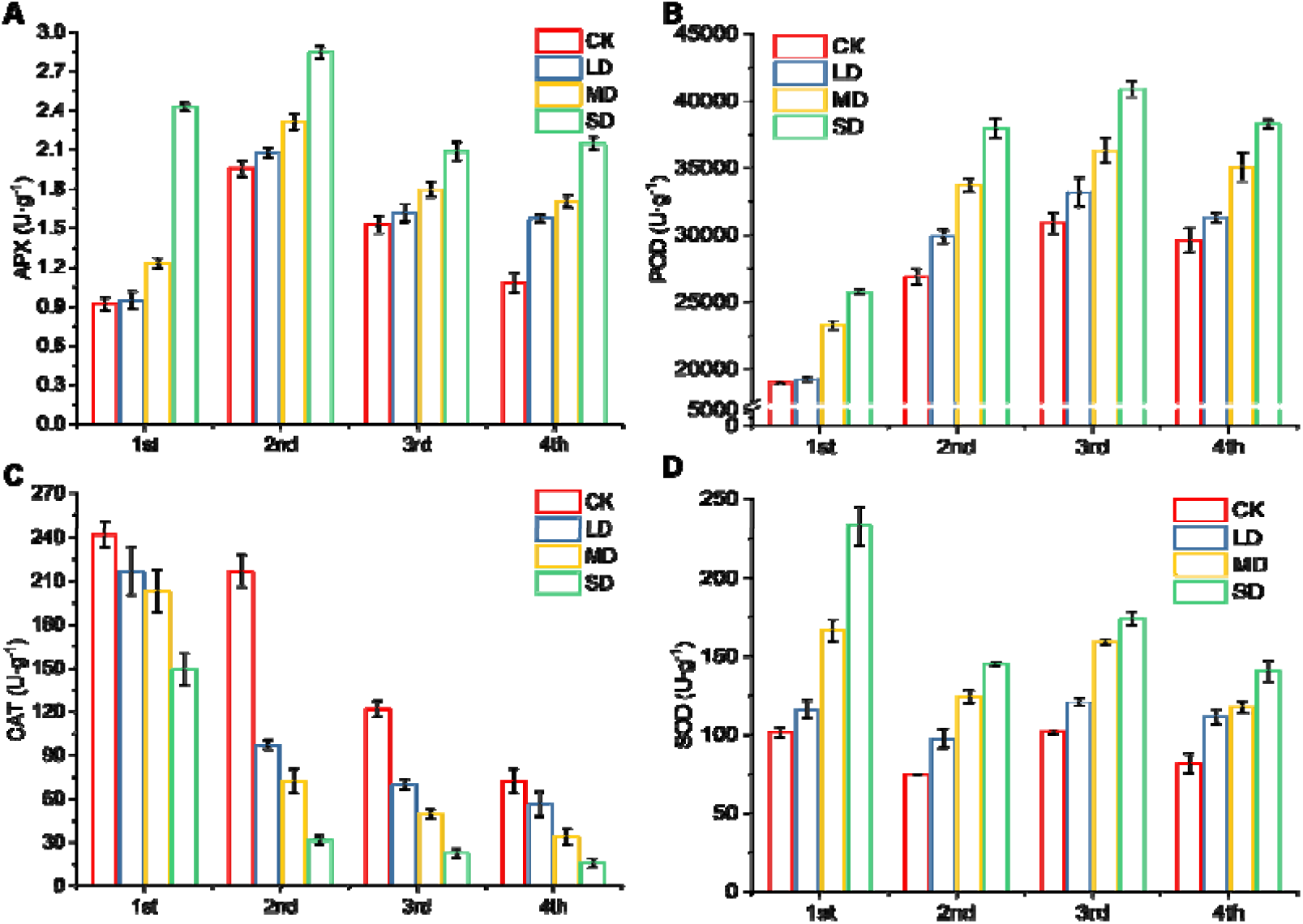
Effect of drought on endogenous protective enzymes in N. tabacum leaves. A. APX; B. POD; C. CAT; D. SOD.

### 3.4. Transcriptome analysis

Principal component analysis (PCA) was used to identify the distinctions among the CK, LD, MD, and SD samples (**Supplementary Fig.2A**). The research revealed that Principal Component 1 explained 83.33% of the variation, while Principal Component 2 contributed 13.01%, resulting in a combined representation of 96.34% of the overall variance. The sample correlation analysis revealed that the correlation coefficients between CK and LD, MD, and SD were 0.74, 0.52, and 0.54, respectively (**Supplementary Fig.2B**). The samples within each DS-treated group clustered closely, suggesting low intragroup variability and high reproducibility.

### 3.5. Differentially expressed genes (DEGs) analysis

To gain a deeper comprehension of the mechanisms by which plants react to drought, we conducted differential expression genes analysis. Comparing with CK, LD treatment resulted in the up-regulation of 5056 genes and the down-regulation of 2427 genes; MD treatment resulted in the up-regulation of 8789 genes and the down-regulation of 6769 genes; SD treatment resulted in the up-regulation of 9072 genes and the down-regulation of 7804 genes. The findings indicated a positive correlation between the severity of drought and the number of genes that were both up-regulated and down-regulated (**Fig. 4A**). After the drought treatment, a total of shared 388 up-regulated genes were identified (**Fig. 4B, C**).

**Fig. 4.**
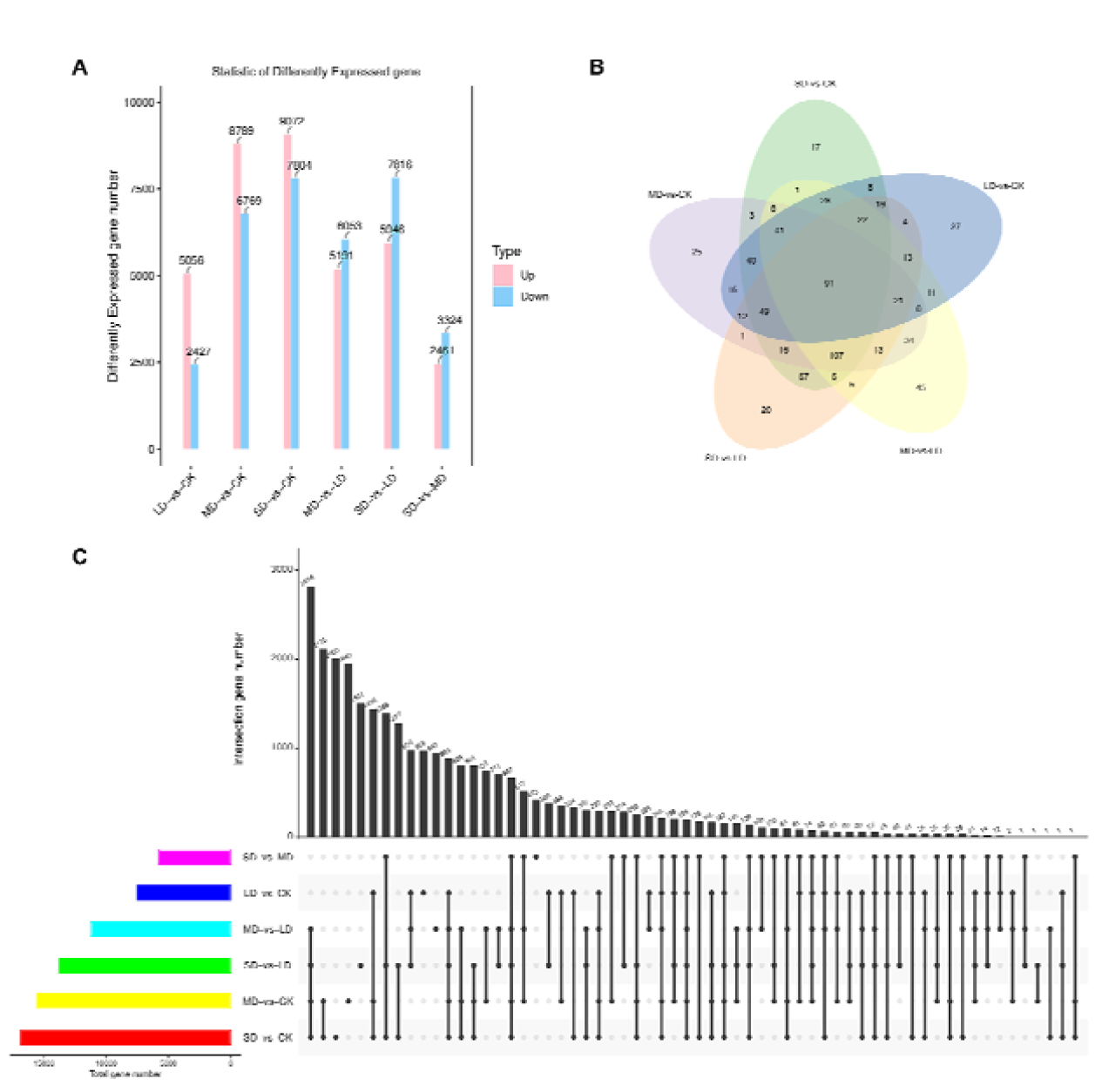
Differentially expressed genes (DEGs) analysis. A, Barplot of DEGs; B, Venn diagram of DEGs; C, UpSet plot of DEGs.

### 3.6. Metabolome analysis

Principal component analysis (PCA) was employed to discern the differences among the CK, LD, MD and SD samples (**Supplementary Fig.3A**). The analysis indicated that Principal Component 1 accounted for 43.9%, while Principal Component 2 contributed 22%, together representing 65.9% of the total variance. The samples within each DS-treated group clustered closely, suggesting low intragroup variability and high reproducibility. Notably, clear separation was observed between the groups, indicating significant differences in the metabolite composition among the plants during the DS treatment (**Supplementary Fig.3A**). Following standardization of the metabolite data, cluster analysis was conducted across all samples (**Supplementary Fig.3B**). The results exhibited distinct clustering among the DS treatment, suggesting notable differences.

### 3.7. Differentially accumulated metabolites (DAMs) analysis

After quantification of all metabolites, the differential metabolites in *N. tabacum* under DS treatment were selected. And then, Top 50 were used for correlation analysis (**Supplementary Fig.4**). Among them, 8 key metabolites related to *N. tabacum* quality were visualized (**Fig.5**). Nicotine, nicotyrine and 3-methylindole showed similar patterns (**Fig.5A, B, C**). Sucrose, isomaltose and fucosyllactose showed similar patterns (**Fig.5D, E, F**). Quercetin 3-Glucosyl-(1->4)-rhamnoside and rutin showed similar patterns (**Fig.5G, H**). The results exhibited distinct patterns among the DS treatments, suggesting notable differences.

**Fig. 5.**
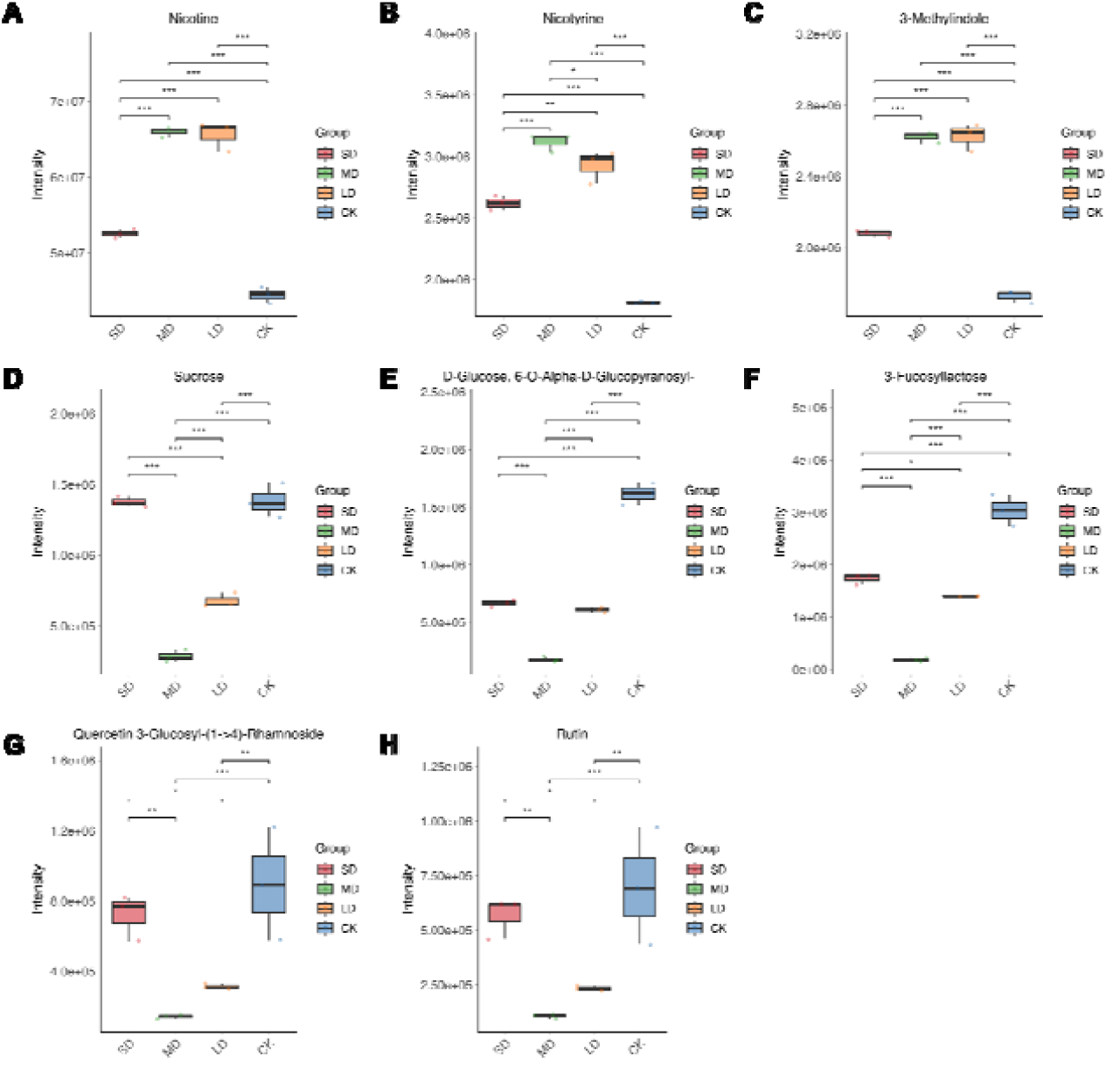
Differentially accumulated metabolites (DAMs) in response to DS in *N. tabacum*.

### 3.8. Quantification of key metabolites in response to DS in *N. tabacum*

The dynamic changes of differential polyphenols under DS treatment have been visualized by the heatmap (**Fig.6**). Through the quantification of four crucial metabolites in *N. tabacum*, it was shown that chlorogenic acid, taxifolin, quercitrin and rutin exhibited a pattern of decrease followed by a subsequent increase. The LD and MD treatments exhibited decreased amounts of these four polyphenols, whereas the CK and SD treatments showed higher levels of these metabolites (**Fig.6**). According to the biosynthetic pathway, all these four polyphenols are originated from phenylalanine, belonging to phenylpropanoids (**Fig. 7A**).

**Fig. 6.**
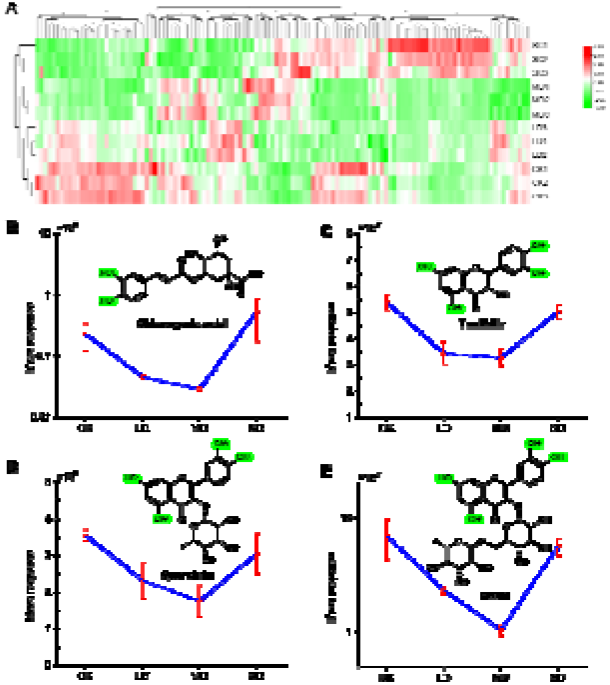
Content of polyphenols in *N. tabacum*. A. Heatmap of polyphenols in *N. tabacum* during DS treatment. B. chlorogenic acid; C. taxifolin; D. quercitrin; E. rutin.

**Fig. 7.**
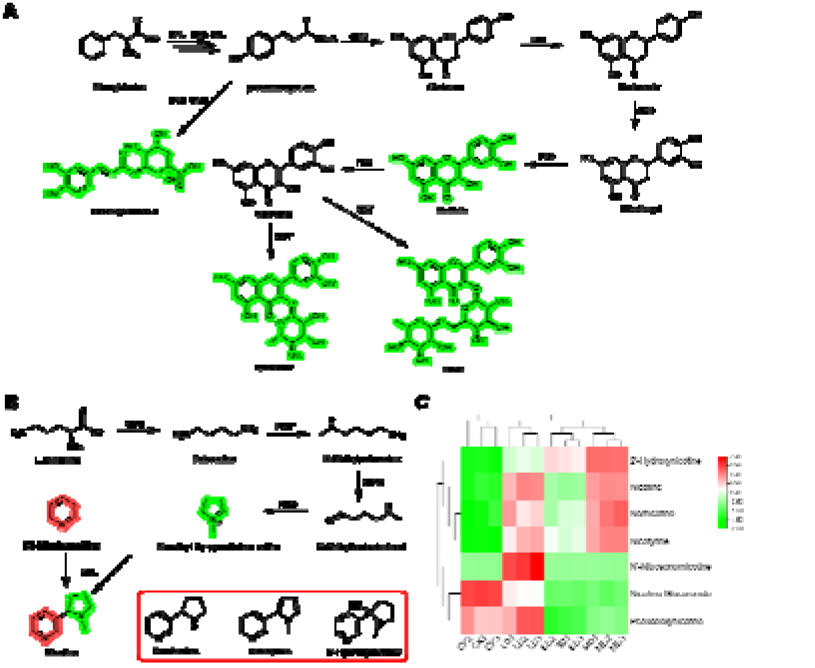
Secondary metabolites in response to drought stress. A, Biosynthetic pathway of polyphenols; B, Biosynthetic pathway of pyridine alkaloids; C, Heatmap of pyridine alkaloids content.

Through the quantification of 7 crucial metabolites in *N. tabacum* (**Fig.7B**), it was shown that both nicotine, nornicotine and nicotyrine exhibited a pattern of increase followed by subsequent decline. The LD and MD treatments exhibited increased amounts of nicotine and nornicotine, whereas the CK and SD treatments showed lower levels of these metabolites (**Fig.7C**).

### 3.9. Combined analysis to identify the pathway in response to DS

To investigate the relationship between metabolites and genes, the KEGG Markup Language (KGML) file in the KEGG database was used to map the network of genes and metabolites. This map will contribute to a more systematic study of the interactions between transcriptome and metabolomics. The KGML results showed the pathways including phenylpropanoid biosynthesis, carbon fixation in photosynthetic organisms, glyoxylate and dicarboxylate metabolism, starch and sucrose metabolism, glycolysis gluconeogenesis and plant hormone signal transduction. Aside from pathways associated with plant development and primary metabolism, the phenylpropanoid biosynthesis pathway, which is the sole secondary metabolic pathway, showed significant enrichment in LD vs CK, MD vs CK, and SD vs CK (**Fig. 8**).

**Fig. 8.**
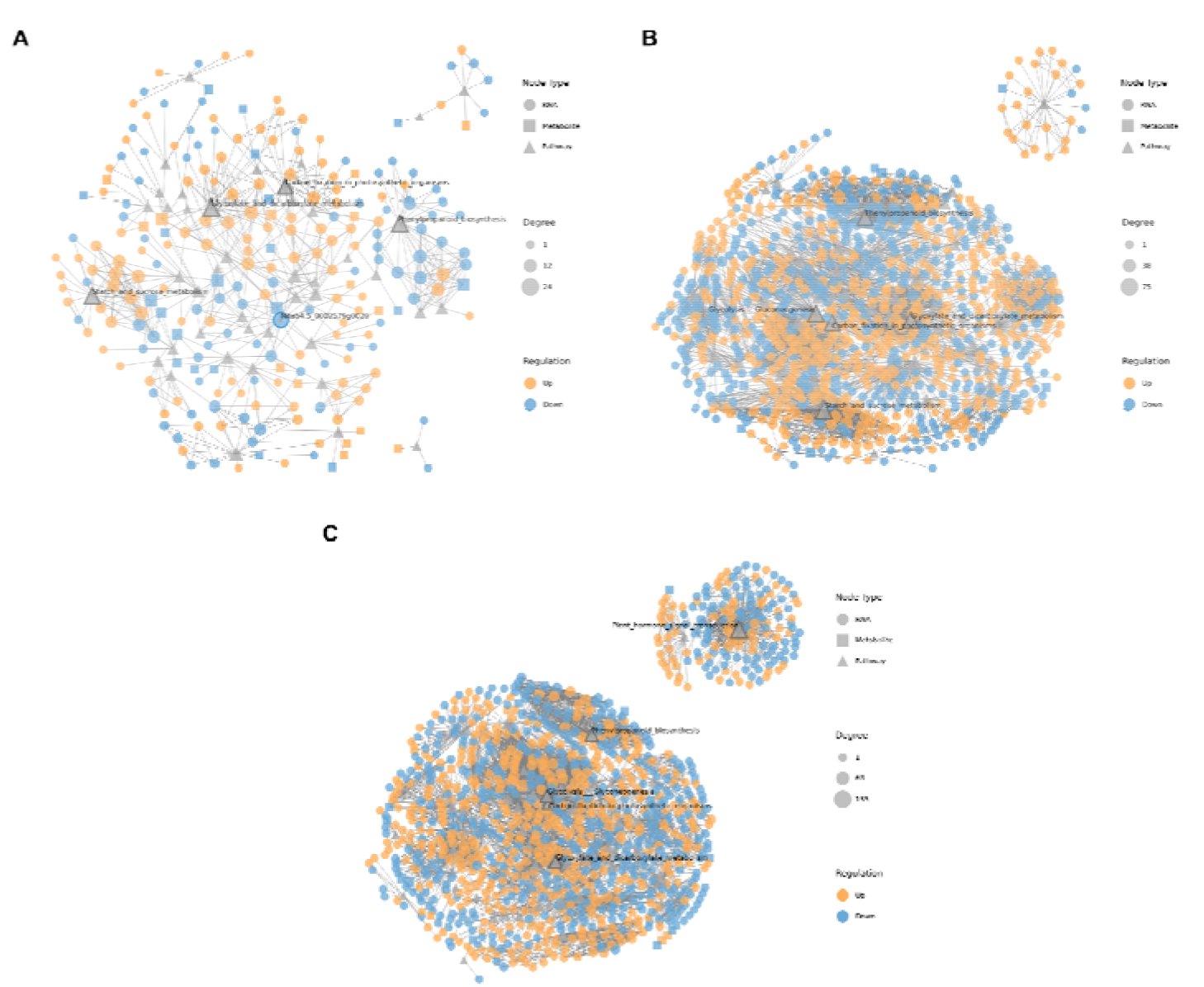
KGML network generated by combined analysis of genes and metabolites. A, LD vs CK; MD vs CK; SD vs CK.

## 4. Discussion

In the face of escalating global environmental challenges, understanding the impact of abiotic stressors on plant physiology is imperative for developing resilient agricultural practices. Fluctuating environmental circumstances, occurring both abruptly and gradually, have an impact on the development and production of plants. Drought stress is the most harmful one among other abiotic stresses with negative impacts on crop growth and development [5, 26, 37]. Global warming and climate change may lead to droughts, resulting in water scarcity that has a detrimental impact on plant growth and production. Hence, plants must see and react to changes in their surroundings and adjust to water shortages in order to survive and thrive. Plants have developed intricate mechanisms to respond to drought stress in order to sustain optimum development under circumstances of water scarcity [38, 39].

### 4.1. Plant growth and development are significantly impacted by drought stress

*N. tabacum* is an important economically crop, plant bioreactor and model plant for biological study grown in more than 120 countries [23, 40]. *N. tabacum* exhibits a significant requirement for water during its reproductive phase, with varying levels of demand observed across different stages of reproduction [26, 41]. When the field’s water holding capacity is below 50%, it has a negative impact on the growth and development of tobacco, leading to a decrease in both yield and quality [42].

Comprehensive investigations have revealed remarkable differences in the phenotype of *N. tabacum* under drought stress [43, 44]. As the severity of drought intensified, observable changes in plant growth, leaf morphology, and overall appearance were noted. In addition, root system development is also influenced a lot by drought stress. These phenotypic variations underscore the profound impact of drought stress on the developmental processes of *N. tabacum* [45]. Drought stress exerts a significant influence on the growth of *N. tabacum*. Reduced water availability impedes essential physiological processes, resulting in stunted growth, decreased biomass accumulation, and compromised plant vigor. The detrimental effects of drought highlight the vulnerability of *N. tabacum* to water scarcity conditions.

In response to drought stress, *N. tabacum* undergoes alterations in the activity of endogenous protective enzymes, such as CAT, POD, SOD and APX [46]. These enzymes play a crucial role in mitigating oxidative damage and enhancing stress tolerance [46-50]. For example, the application of zinc-oxide nanoparticles to tomato plants resulted in oxidative stress, which was characterized by increased MDA content and accumulation of H_2_O_2_, as well as increased activities of antioxidant enzymes, such as SOD, CAT, APX [51, 52]. Additionally, the cotton’s catalase (CAT) activity exhibited a decline in response to drought stress [53]. The observed changes in enzyme activity reflect the plant’s adaptive strategies to cope with drought-induced oxidative stress and maintain cellular homeostasis.

### 4.2. Phenylpropanoids mediated drought tolerance in *N. tabacum*

Phenylpropanoids are secondary metabolites important for plant development. They play essential roles in different physiological processes including photosynthetic activity, hormonal regulation, nutrient mineralization, metabolism and reproduction [54]. Plant phenylpropanoids are also vital for the resistance to abiotic stresses [55, 56]. In their living habitat, plants are constantly exposed to several constraints such as heavy metals, cold, heat, drought, ultraviolet radiation, salinity, etc., which are detrimental to their productivity. To survive, plants develop tolerance to the stresses, and this evolution process results in the accumulation of phenylpropanoids in different tissues as a response to the harmful conditions [57]. Phenylpropanoids are composed of different compounds including flavonoids, coumarins, monolignols, stilbenes and various phenolic acids [58, 59]. Resveratrol is a natural phytoalexin produced by plants in response to biotic and abiotic stresses [60]. The content of rutin and quercetin increased in buckwheat with moderate drought stress. Rutin content ranged from 16.24 to 32.5 mg/g dry weight. Besides, phenolic, flavonoid, and rutin content are correlated with higher antioxidant properties [61]. Chlorogenic acid (CGA) belongs to a class of secondary metabolic products involved in transcriptional activation of the phenylpropanoid pathway under pathogen infection and abiotic stress events [62]. It has been demonstrated that taxifolin functions as a stimulator that recruit beneficial plant bacteria to fight biotic stressors like pathogenic bacteria [63].

Collectively, the integration of plant phenotypes, physiological parameters, transcriptome, and metabolome studies has elucidated the involvement of polyphenols in the response of *N. tabacum* to drought stress. Through mining and identification of superior stress resistance candidate genes, we can get further understanding and knowledge for the future breeding of *N. tabacum* germplasms with enhanced resistance to abiotic stress.

## 5. Conclusions

This study investigated the plant phenotypes, defensive enzyme activities, and specific metabolites in *N. tabacum* in relation to drought stress. This work focused on examining the response of *N. tabacum* to varying levels and durations of drought stress. The aim was to gain insights into the mechanism by which *N. tabacum* responds to drought stress and establish a theoretical foundation. Thorough investigations into how *N. tabacum* responds to drought stress have yielded useful knowledge about the phenotype, physiological adjustments, genetic control, and accumulation of substances employed by *N. tabacum* to cope with water scarcity. Understanding these adaptive responses is crucial for developing strategies to enhance drought tolerance and resilience in *N. tabacum*, thereby ensuring sustainable crop productivity in arid and water-limited environments under global warming.

## CRediT authorship contribution statement

Conceived and designed the experiments: MY, QY. Performed the experiments: ZF, ZR, HW, DW. Analyzed the data: MY, ZF HW, DW. Contributed reagents/materials/analysis tools: MY, ZF, ZR HW, DW. Wrote the paper: MY, ZF, AJ. All the authors contributed to editing the manuscript.

## Acknowledgments

We appreciate financial support from the Key Technology R&D Program of Henan Province (242102110240, 232102110053), the Special Support Fund for High-level Talents and skills improvement of Henan Agricultural University (30501474), Luoyang Tobacco Company Scientific and Technology Program (2019410300270141), China National Tobacco Company-ShaanXi Provincial Company Scientific and Technology Program (2022610700260183; 2021611000270042).

## Data availability

Data will be made available on request.

